# Linking Cyanobacterial Genomes to Toxin Dynamics Through Genome-Resolved Metagenomics

**DOI:** 10.64898/2026.02.24.707707

**Authors:** Autumn Pereira, Fernando Martínez-Jerónimo, David P. Fewer, Dana F. Simon, Miriam Hernández-Zamora, Laura Martínez-Jerónimo, Paloma Antuna-González, Gabriel Munoz, Sébastien Sauvé, B. Jesse Shapiro, Nicolas Tromas

## Abstract

Global climate change and nutrient pollution from agricultural systems increase cyanobacterial bloom and toxin release events, and will pose a significant concern for public health and safety over the coming decades. Many *in situ* studies have focused on environmental, chemical, and microbial community changes and their impact on cyanobacterial bloom frequency and toxicity. However, fine-scale genetic differences in the genomes of bloom-forming cyanobacteria may also impact the quantity and types of toxins produced. Metagenomic approaches allow resolution of strain- and nucleotide-level changes within microbial communities and can improve our understanding of the factors that affect cyanobacterial bloom dynamics and toxicity. Here, we conduct a metagenomic analysis of the bloom-forming cyanobacterial genus *Microcystis* across a 10-month lake time series from the Valle de Bravo Reservoir, to assess how within-genus genotype-level changes are linked to intracellular toxin production and extracellular toxin release, as well as how single nucleotide variation may affect the types of microcystin toxins produced. Our results demonstrate that the abundances of both toxigenic and non-toxigenic *Microcystis* genotypes are significantly related to microcystin toxin concentrations. In addition to these genome-wide (”strain”-level) associations, specific single nucleotide variants show strong associations with chemical variants of microcystin in the environment. Our work highlights the importance of fine-scale analysis of microbial community composition for understanding cyanobacterial bloom dynamics and toxin production.

## Introduction

Cyanobacterial blooms have been increasing in frequency and duration since the onset of the industrial revolution, in a phenomenon directly linked to anthropogenic-induced climate and land use changes [1, 2]. These blooms can have negative impacts through declining lake health, disruption of food web dynamics, and notably, the release of harmful cyanotoxins [3, 4]. Much effort has been made to develop predictive models for cyanobacterial bloom formation and toxin release [5]. However, current models based on abiotic lake parameters and are limited in their ability to predict bloom formation at-scale [5]. It has become increasingly clear that incorporating biological factors and genomic characteristics of the cyanobacteria involved in these toxin production and release events is an important step in improving our models and strategies for bloom control [5, 6].

Cyanotoxins pose a significant risk to public health, as they can have severe impacts on the digestive and nervous systems [1]. The hepatotoxin microcystin is the most well-studied of the eight major classes of cyanotoxins, due to its ubiquity in association with harmful cyanobacterial blooms [7]. There are more than 270 known chemical variants (or congeners) of microcystin, varying significantly in their toxicity [8]. Most of the chemical variation of microcystin congeners results from amino acids substitutions at positions 2 and 4, or demethylation at positions 3 and 7 in the cyclic peptide [8]. Microcystins can be synthesized by numerous cyanobacteria, including the bloom-forming genera *Microcystis*, *Anabaena*, *Dolichospermum* and *Planktothrix* [9]. The toxin is synthesized intracellularly using non-ribosomal peptide synthetases and polyketide synthase enzyme complexes encoded in microcystin *mcy* biosynthetic gene clusters, which canonically consist of ten genes (*mcyA-J*) organized into two operons, with expression driven from a single bidirectional promoter [9]. However, the number of biosynthetic genes, gene order, and promoter location can vary between producing organisms [10, 11]. The exact microcystin congeners produced in a given cyanobacterium varies due to genetic modifications, including nucleotide substitutions, deletions, and large recombination events that affect the substrate specificity and function of microcystin biosynthetic enzymes [12, 13, 9]. Once produced, microcystin is released to the environment primarily through the processes of senescence and cell lysis [3].

The reasons why cyanobacteria produce cyanotoxins are not well understood. Leading hypotheses to rationalize cyanotoxin production include response to abiotic factors and nutrient sources, competition and predator deterrence, and bloom formation and maintenance [14, 15]. Additionally, the dynamics of potential producers and non-producers, and the factors that shape the abundances of these two groups relative to one another, remain unclear. Toxin-production ability is typically assigned at the genotype-level due to significant variation in microcystin synthesis ability within a given species [16]. Therefore, traditional amplicon-based approaches to microbial community analysis are limited in their ability to identify meaningful variation in microbial communities associated with toxin production. Numerous studies tracking the presence of *mcy* marker genes using quantitative PCR have been carried out to quantify the abundance of potential microcystin producers (e.g. [17]). However, some key studies have shown that the abundances of *mcy* marker genes do not always correlate with microcystin toxin concentration [18].

Recent untargeted metagenomic studies have provided new insights into the activity and composition of *Microcystis* populations in lakes, including the prevalence and expression of *mcy* biosynthetic gene clusters with truncated genes [10]. These approaches are powerful because they provide insights into the structure and activity of these microbial communities in much finer detail than is possible using targeted amplicon sequencing approaches.

Here, we use genome-resolved metagenomics to analyze *Microcystis* communities in the Valle de Bravo reservoir (Mexico), to yield fine-scale resolution of the genomic changes associated with microcystin toxin production. We used this approach to assess how the abundances of various *Microcystis* genotypes and single nucleotide variation contribute to the microcystin toxin production and release in this freshwater system, as well as the links between *Microcystis* genotype abundances and environmental variation. Our dataset comprised a 10-month time series in a watershed with constant bloom presence, where *Microcystis* was the only potential microcystin-producing genus observed [19]. We hypothesized that the dynamics of microcystin congeners observed in the reservoir can be explained in part by both the ratios of toxigenic and non-toxigenic *Microcystis* genotypes, as well as the genetic variation within the microcystin synthetase biosynthetic gene cluster. Our analyses demonstrate that non-toxigenic genotypes coexist with toxigenic genotypes, throughout a range of microcystin toxin concentrations, and that the abundances of both toxigenic and non-toxigenic genotypes are significantly associated with intracellular and extracellular toxin concentrations observed in the reservoir. We also identified numerous nucleotide positions within the microcystin synthetase biosynthetic gene cluster which are associated with the production of non-dominant microcystin congeners, suggesting that small genetic variation in the *mcy* biosynthetic gene cluster can have significant impacts on the chemical variants of microcystin produced in natural environments. This work highlights the utility of metagenomic approaches for analysis and prediction of cyanobacterial bloom dynamics and cyanotoxin release, and contributes to the increasing evidence that toxigenic and non-toxigenic *Microcystis* genotypes coexist in nature.

## Methods

### Sampling

The Valle de Bravo reservoir is a highly eutrophic, monomictic lake located near Mexico City, Mexico (19°21’30” N, 100°11’00” W) [20]. This reservoir has a surface area of 18.6 km^2^ and is the largest of the seven reservoirs in the Sistema Cutzamala which provide 30% of the drinking water supply for more than 26 million inhabitants of the Mexico City Conurbation [21]. The region has a subhumid climate, a dry season spanning from November to May, and a wet season from June to October [20].

Sample collection and water chemical analysis are detailed in [19]. Briefly, duplicate water samples were taken once per month over a 10-month period in 2019 from 6 distinct locations in the watershed. These locations were chosen to capture the influence of the nearby tributaries connected to the lake, as well as the influence of anthropogenic activities along the coastline.

Samples were vacuum-filtered within 6 hours following sample collection to capture cyanobacterial biomass, and subsequently frozen at −80°C to preserve cells until processing. Intracellular and extracellular concentrations of 10 common microcystin congeners were assessed using Lemieux von Rudloff oxidation followed by solid phase extraction and ultra-high-performance liquid chromatography tandem mass spectrometry, as detailed by [22]. Further descriptions of sample processing are described in [19].

### DNA Extraction and Sequencing

An aliquot of 50-250 mL of each water sample, depending on the biomass, was filtered through a 0.22 *µ*m nitrocellulose filter membrane prior to DNA extraction. Filters were kept at –20°C until analysis. DNA extraction was performed using the DNeasy PowerWater Kit (Qiagen) following manufacturer instructions. DNA was subsequently quantified with a Qubit v.2.0 fluorometer and sequenced with Genome Quebec sequencing platform. Library preparation was completed using the NEB Ultra II kit (New England Bio Labs) and sequenced with NovaSeq 6000 S4, (2x 150bp; Illumina Inc. San Diego, CA, USA).

### Bioinformatic Analyses

#### Generation of metagenome-assembled-genomes

We began by using fastp (v0.20.1, [23]) to trim adapter sequences and remove low-quality reads. The resulting cleaned reads were then co-assembled by lake using Megahit (v.1.2.9, [24]). Metabat2 (v2.14, [25]) and Vamb (v3.0.8, [26]) were then used to bin contigs, and these contigs were subsequently dereplicated into metagenome-assembled genomes (MAGs) using DAS Tool (v1.1.6, [27]) as previously described [19]. MAGs were then screened for contamination, and affected genomes were removed using MAGpurify v1.0 with phylo-marker, GC-content, and tetranucleotide frequency modules [28]. We used CheckM (v1.2.2, [29]) and metashot prok-quality (v1.3.1, [30]) to evaluate the recovered MAGs and retained only those that met MIMAG criteria [31]. Taxonomy was assigned to each MAG using GTDB-Tk (v.1.5.0, data release 207 v2, [32]) with default settings. MAGs were then dereplicated using dRep (v3.4.3, [33]) with default parameters. We finally used prokka (v4.0, [34]) using default settings on filtered MAGs (completeness*>*75%, contamination *<*10%) for gene annotation. A total of 17 *Microcystis* MAGS were recovered from the metagenome assembly process, which was reduced to 12 MAGs based on the CheckM quality check step (Table S1). As *Microcystis* was found to be the only bacterial genus responsible for microcystin toxin production in our study system [19], we conducted downstream analyses only on this set of *Microcystis* MAGs.

### Reference *Microcystis* Database Creation

We complemented our *de novo* assembly approach with a reference-based assembly to ensure that we captured the maximum amount of genetic diversity in our analyses. We conducted identical control analyses in parallel using a set of public *Microcystis* reference genomes [35]. We used CheckM v.1.2.2 [29] to quantify the completeness and contamination of the reference genomes and removed any with completeness *→* 90% and contamination *↑* 10%. This resulted in a set of 330 unique genomes, which was reduced to 310 after filtering.

### Phylogenetic Relationships

We first used prokka to annotate the reference genomes and the MAGs assembled from our environmental samples (v.1.14.5, [34]). Following the annotation step, we used ultrasensitive Diamond BLASTP (v.2.0.15, [36]) to search the reference genomes and MAGs to identify genes belonging to the *mcy* biosynthetic gene cluster. We constructed a dataset of *mcy* biosynthetic gene sequences as previously described [16]. We identified “hits” as sequences in the genomes of our database with *↑* 30% query cover and *↑* 60 % identity compared to the *mcy* biosynthetic genes in *mcy* gene data set. We further limited the maximum number of target sequences to 1. Toxigenic genotypes were defined as those with all ten of the *mcy* biosynthetic genes, which compose complete *mcy* biosynthetic gene cluster in genotypes of the genus *Microcystis*. We classified non-toxigenic genotypes as those with at most one of the core *mcy* biosynthetic genes (*mcy* A-E, *mcy* G). Genomes that contained at least two of the core genes required for microcystin synthesis were classified as “ambiguous”, given that microcystin synthesis may be possible with a partial *mcy* biosynthetic gene cluster [10]. We then used these annotated MAGs and reference genomes to construct a pangenome and identified sets of core and accessory genes using the MAFFT tool in Roary (v.1.7.7, [37]). Core genes were defined as those which had blastp *↑* 0.9, were present in *↑* 99% of the genomes in our database, and the maximum number of clusters set to 70 000. The resulting alignments contained 238 core (99-100% of genomes) and 780 soft core genes (95-99% of genomes). We identified phylogenetic relationships among the genomes in our database using IQ-TREE2’s extended model selection and tree inference tools, constructing and comparing phylogenetic trees with 1,000 ultrafast bootstrap replicates [38, 39, 40]. Visualization of the most parsimonious unrooted tree identified by IQ-TREE was achieved using iTOL (v.7, [41]). Based on visual assessment, two genomes were removed from the final phylogenetic tree, given significant distance of these two genotypes relative to the rest of the *Microcystis* genomes downloaded from NCBI.

### Analysis of Sample Community Composition

We ensured that the sample read depth did not affect the ability for genotypes to be detected between samples by downsampling our sample metagenome reads using seqtk (v.1.3, [42]). Samples were downsampled to 18 million reads, and two samples, the October sample for the second site and the July sample for the sixth site, were removed from our analyses due to low sample reads (*<* 5 million). We used StrainGE (v.1.3.7 [43]) to assess the genotype-level variation in our metagenome samples. We conducted two separate analyses, the first using our reference database, and the second analysis using only our *Microcystis* MAGs isolated as described above. A k-mer set was created from the genome sequences for each genome in our two databases, and genomes with high Jaccard similarity (*>* 0.90) were grouped together. We used StrainGE to map our downsampled metagenome reads to the genomes in our databases to identify the genotype-level composition of our samples. Plots of genotype-level composition of *Microcystis* communities were created in RStudio using the ggplot2 package [44, 45].

We assessed the relationship between the abundances of toxigenic *Microcystis* MAGs and microcystin concentrations, by conducting linear regression using the ratio of the abundances of toxigenic to non-toxigenic *Microcystis* MAGs as a predictor and log-transformed microcystin concentration as the response variable.

Additionally, we conducted multiple linear regression to identify the relationships between *Microcystis* MAGs and total toxin concentration. A redundant model was generated which included the Hellinger-transformed abundances of all *Microcystis* MAGs as predictors and the log-transformed sum of the intracellular or extracellular toxin concentrations as the response variable. Backwards model selection based on AIC was carried out to determine the best fitting model using the vegan package [46]. All linear regression analyses were conducted in R [45].

Finally, a redundancy analysis was conducted to identify specific MAGs which may be associated with the abundances of particular microcystin congeners. Redundancy analysis was carried out using the vegan package in R, with a response matrix of log-transformed intracellular or extracellular toxin concentrations, and a predictor matrix of Hellinger-transformed MAG abundances. The best model was selected using backwards model selection based on AIC using the vegan package in R [46].

### Analysis of Genetic Variation of in the *mcy* operon

We assessed the relationship between single nucleotide variation in the microcystin synthetase operon and the abundance of microcystin congeners to determine whether genetic variation explains congener diversity in our study system. We began by analyzing the relationships between the *mcy* biosynthetic gene clusters of our MAGs and a reference *mcy* biosynthetic gene cluster from *Microcystis aeruginosa PCC 7806SL* (NCBI Accession no. *GCA* 041506625.1). We used ANTISMASH [47] to identify *mcy* biosynthetic genes in our MAGs and reference genomes. We then extracted the region containing the *mcyA, mcyB* and *mcyC* genes in the *mcy* biosynthetic gene cluster from the ANTISMASH output and aligned and visualized the region using Digialign [48].

The *mcyABC* region of the *mcy* biosynthetic gene clusters extracted from each of our MAGs were nearly identical in gene order and sequence composition. We therefore conducted further analysis to investigate if single nucleotide variation in the *mcyABC* region observed in our samples could explain the abundances of different toxin congeners.

We aligned the downsampled metagenome reads from each sample to the sequence of the *mcy* biosynthetic gene cluster extracted from *Microcystis aeruginosa PCC 7806SL* using Bowtie2 (v.2.5.1, [49]). This alignment was then analyzed using InStrain (v1.7, [50]) to identify variable nucleotide positions in the sample metagenomes relative to the reference. Only variable bases with at least 5x coverage were retained in our analyses. We then extracted the frequency of observing single nucleotide variation at each position in the *mcyABC* region for further statistical analysis.

We conducted redundancy analysis using the frequency of single nucleotide variation (SNV) at different sites in the *mcyABC* region as predictors and the concentrations of toxin congeners as the response matrix to identify links between *mcy* biosynthetic gene nucleotide sequences and the production of various toxin congeners. As redundancy analysis does not allow the inclusion of NA values in the response or predictor matrices, we removed 11 samples which had greater than 200 NA counts across the *mcyABC* region. We then removed any positions with low variation (identified as positions with *<* 4 unique values for SNV frequency observed across all samples) and finally removed all positions which had a value of NA for at least one of the sample metagenomes. We ran redundancy analysis with the resulting set of 26 positions within *mcyABC* in 41 sample metagenomes, and identified the best fitting model based on AIC using backwards model selection [46].

Lastly, we assessed the utility of MAG-based and SNV-based approaches in understanding microcystin toxin congener abundances by comparing the optimized redundancy models containing MAG-only and SNV-only predictors to a model containing the MAG and SNV predictors from both optimized models. For these comparisons, we re-ran the redundancy models using only the 41 sample metagenomes identified above to be suitable for SNV analysis. We then compared the adjusted *R*^2^ values from these three models, as well as the significance of factors between each individual model and the combined model with both SNV and MAG predictors.

## Results and Discussion

### Closely related *Microcystis* MAGs vary in their ability to produce microcystins

We previously identified 17 *Microcystis* MAGs in the Valle de Bravo Reservoir [19]. Quality filtering based on completeness and contamination reduced this number to 12 MAGs, which clustered in two locations in our phylogenetic tree (Figure 1).

**Figure 1:**
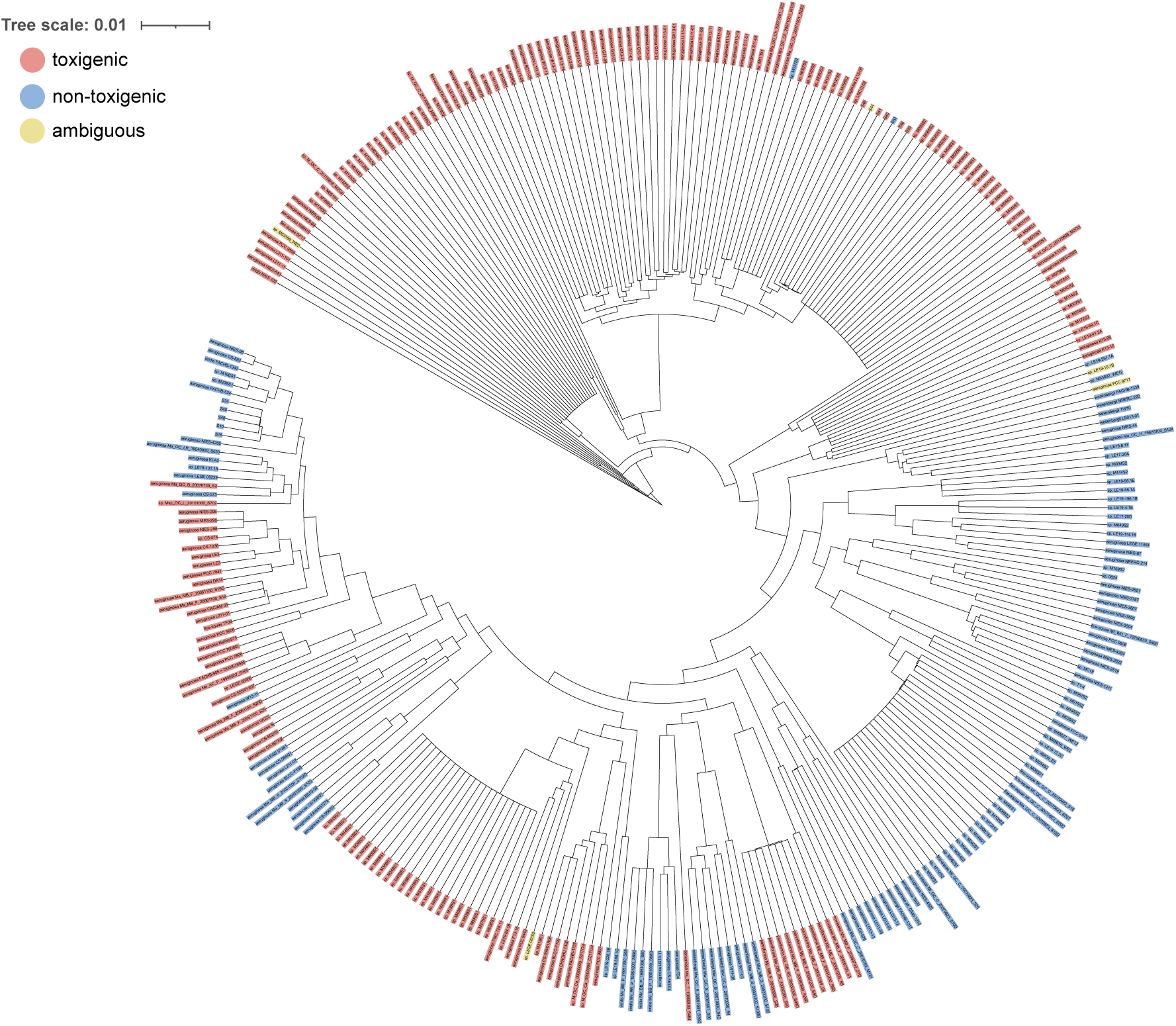
Phylogenetic tree of *Microcystis*MAGs and reference genomes used in this study. The tree includes 308 of the 330 *Microcystis* genomes available in NCBI as of March 27, 2024. Genomes identified as toxigenic (encoding the full set of *mcy* biosynthetic genes) are shown in red, while non-toxigenic genomes (encoding at most one *mcy* biosynthetic gene) are shown in blue. Genomes which are ambiguous in their ability to synthesize microcystin, containing at least 2 of the core *mcy* biosynthetic genes, are shown in yellow.

Our phylogenetic tree consisted of 308 *Microcystis* reference genomes and 12 *Microcystis* MAGs, of which 192 (60.0%) genomes encoded all ten *mcy* biosynthetic genes, 123 genomes (38.4%) encoded 1 or fewer core *mcy* biosynthetic genes (*mcyA-E* and *mcyG*), and 5 (1.6%) genomes contained 2 or more core *mcy* biosynthetic genes (Figure 1). The tree topology did not show clustering based on toxin-producing identity; this result is unsurprising, since previous studies have shown that although microcystin toxin production appears to be an inherited trait rather than transferred horizontally, numerous loss events have led to a sporadic distribution of the *mcy* biosynthetic gene cluster among genomes of the same genus [51].

MAGs that encoded the full set of *mcy* biosynthetic genes clustered with toxigenic *Microcystis aeruginosa LL13-06* and non-toxigenic *Microcystis sp. LSC13-02*, while non-toxigenic genotypes clustered only with non-toxigenic references. These results, which suggest that there are two main *Microcystis* species in our set of recovered *Microcystis* MAGs, are supported by FastANI outputs comparing only the *Microcystis* MAGs to one another. These comparisons show two separate clusters with ANI *<* 95%, consisting of non-toxigenic and toxigenic MAGs with ANI *>* 97% (Figure 2).

**Figure 2:**
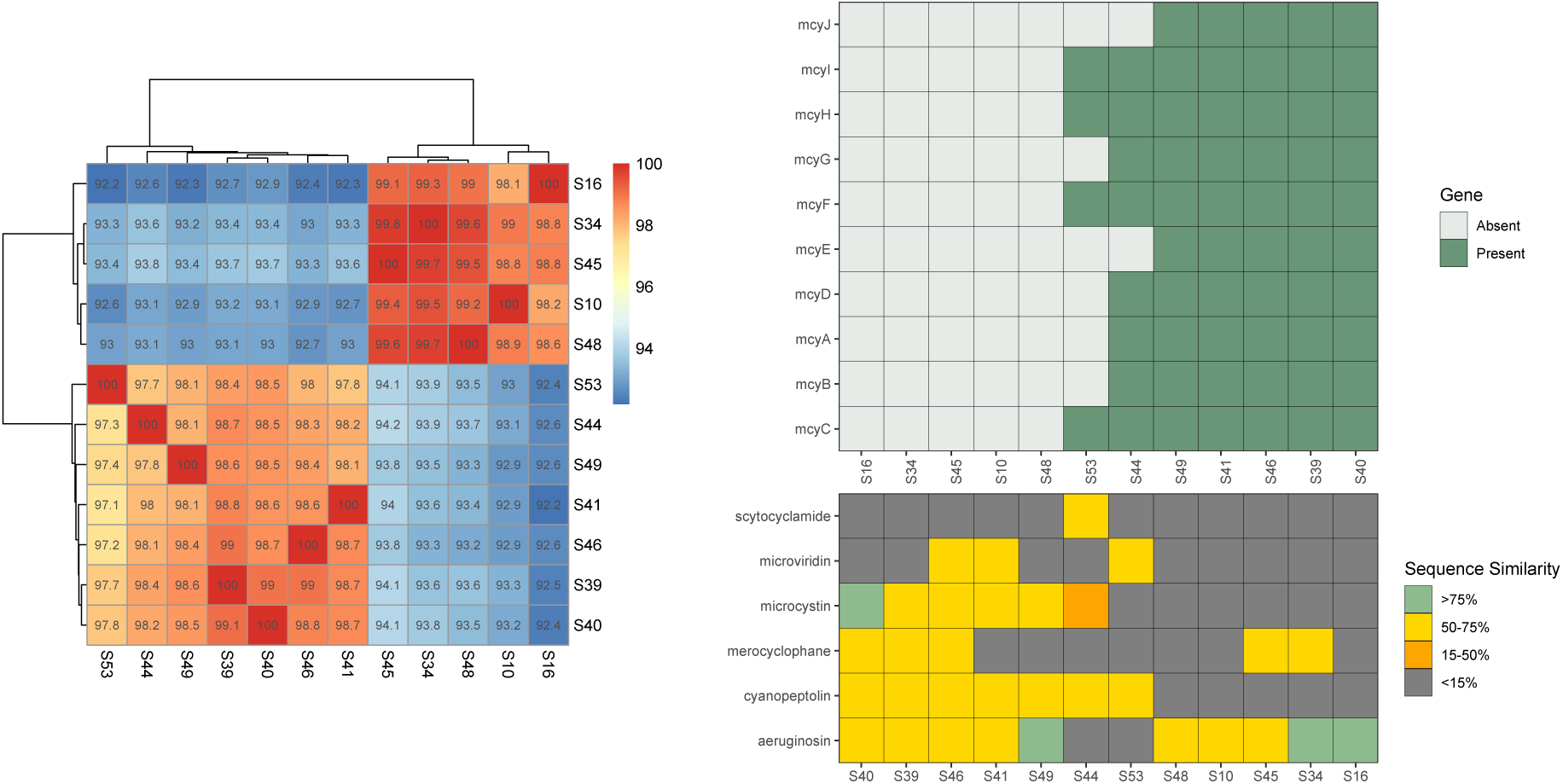
Two genetic clusters of MAGs differ in their toxin gene repertoires. (left) Heatmap of ANI values and clustering of the MAGs included in our study. ANI values were assigned using FastANI, and showed two distinct clusters of MAGs with *>* 97% similarity. (upper right) Heatmap of the *mcy* genes identified in our MAGs. Genes were considered present if they were identified in the recovered MAG genomes at *↑* 30% query cover and *↑* 60% identity. While most MAGs contain either all or none of the *mcy* biosynthetic genes, S44 contains all but one (*mcyE*) of the core *mcy* genes, while S53 contains only one of the core *mcy* genes. (lower right) Heatmap of the sequence similarity output by ANTISMASH of additional biosynthetic gene clusters identified in our MAGs. While some MAGs do not contain the *mcy* biosynthetic gene cluster, all MAGs contained biosynthetic gene clusters which could produce other natural products, notably aeruginosin.

### Toxigenic and non-toxigenic *Microcystis* genotypes coexist through a range of microcystin concentrations

Genotype-level analysis of the *Microcystis* community composition in our samples using MAG- and reference-based approaches both demonstrated that toxigenic genotypes of *Microcystis* coexist with non-toxigenic genotypes among a wide range of intracellular microcystin concentrations and timepoints (Figures 3, S4). Interestingly, we also observed an increase in the concentrations of intracellular microcystin toxins during the months of June through September at all sites, while the abundance of potential producers during these timepoints remained similar to or lower than the relative abundances observed during months with low microcystin abundances (Figures 3, S4) as well as low abundances of *Microcystis* MAG kmers relative to all kmers from the DNA in our samples (Figure S1) at this timepoint at all sites.

**Figure 3:**
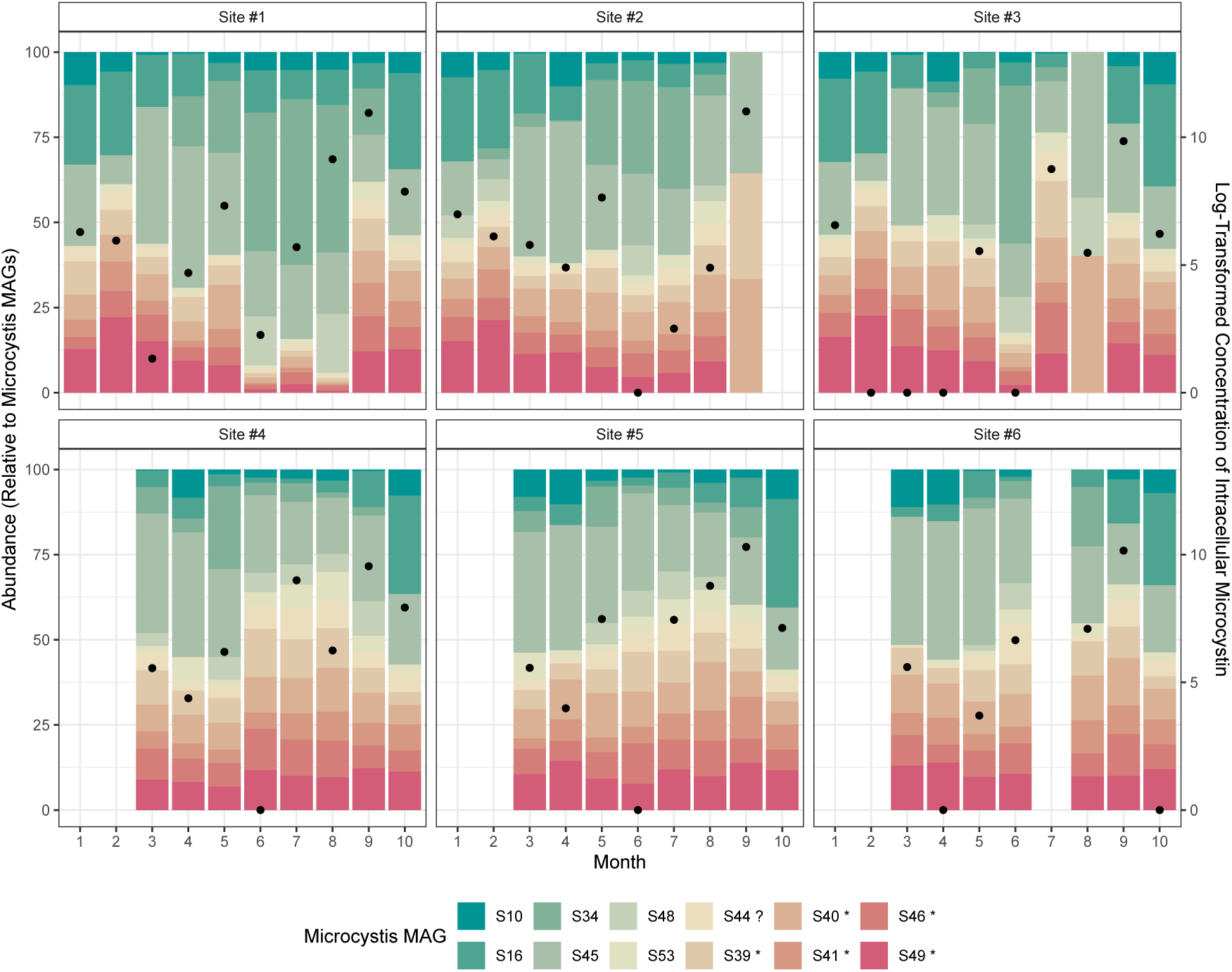
Toxigenic and non-toxigenic MAGs coexist over space and time. Relative abundances of *Microcystis* MAGs at different timepoints and locations in the Valle de Bravo Reservoir. The left y-axis corresponds to the stacked bar plot, where the numerical value represents the relative abundance of the MAG in the *Microcystis* community, with MAG IDs included in the legend. MAG IDs with an asterisk beside the name denote toxigenic MAGs, while those with a question mark represent genomes which are ambiguous in their ability to synthesize microcystin. The right y-axis and points represent the log-transformed sum of the concentrations of the 10 measured intracellular microcystin toxin congeners. Toxigenic and non-toxigenic MAGs co-occurred at all sites and timepoints in our study, and high abundances of microcystin were observed at all sites in June through September. However, increases in toxin concentrations were not always coupled with high abundances of toxigenic MAGs.

These results align well with other *in situ* work using metagenomic sequencing [10] and DGGE [52] approaches to quantify *Microcystis* community dynamics in freshwater systems, which have shown similar results of coexistence of non-toxigenic and toxigenic genotypes. Additionally, some *in vitro* studies have shown that microcystin synthesis ability itself is insufficient to predict the outcome of competition between *Microcystis* genotypes, and that other factors such as nutrient and light availability represent important predictors of the outcomes of competition [53, 52]. Given that these environmental factors can be highly spatially and temporally variable in lake systems, the results of these laboratory studies may provide some insight into possible explanations for the coexistence of toxigenic and non-toxigenic *Microcystis* genotypes.

Moreover, these results may suggest that, for closely related species of *Microcystis*, microcystin production may be less important for allelopathic purposes and generally does not result in competitive exclusion in nature. However, our analyses do not completely rule out the possibility for microcystin synthesis as an allelopathic or competitive strategy. While most of our MAGs contained genes required for synthesizing other allelopathic chemicals, such as aeruginosin, our study did not quantify the abundance of all of these possible allelopathic compounds. It is therefore possible that the genotypes we classified as non-toxigenic based on microcystin synthesis ability were able to compete based on their ability to synthesize aeruginosin or other allelopathic compounds. While the ability for *Microcystis* species to synthesize aeruginosin has been recognized in other work, the competitive ability of cyanobacterial species producing different toxins is an underrepresented subject in the literature. Future studies should consider analysis of a wider panel of cyanotoxins *in situ* to more accurately assess the competitive ability of *Microcystis* genotypes which do not synthesize microcystin but might produce other allelopathic compounds.

### The relative abundance of *Microcystis* MAGs is associated with intracellular and extracellular microcystin concentrations

Several microcystin variants were present and co-abundant in Valle de Bravo as previously described [19]. *Microcystis* was identified as the only genus carrying the *mcy* biosynthetic gene cluster in this system, and five of our 12 MAGs contained the full *mcy* biosynthetic gene cluster, allowing them to potentially produce microcystins. Given the high proportion of recovered *Microcystis* MAGs with microcystin synthesis abilities, we conducted regression and redundancy analyses to identify links between microcystin concentrations and MAG abundances.

We found that intracellular, but not extracellular, microcystin concentrations are significantly correlated with the ratio of toxigenic to non-toxigenic MAG abundances (Figure 4). This result may be explained by the different processes leading to the observation of intracellular and extracellular toxins. While intracellular toxins are synthesized by living cells and may therefore be more reflective of active members of the *Microcystis* community, extracellular toxins are typically observed after cell senescence and lysis. Therefore, extra-cellular toxin concentrations may be more strongly associated with community turnover, and therefore the lack of significant relationship between the ratio of toxigenic to non-toxigenic MAGs and extracellular toxin concentration may be because toxigenic *Microcystis* might not be abundant at timepoints with high extracellular microcystin, since the cells which the microcystin came from have been lysed.

**Figure 4:**
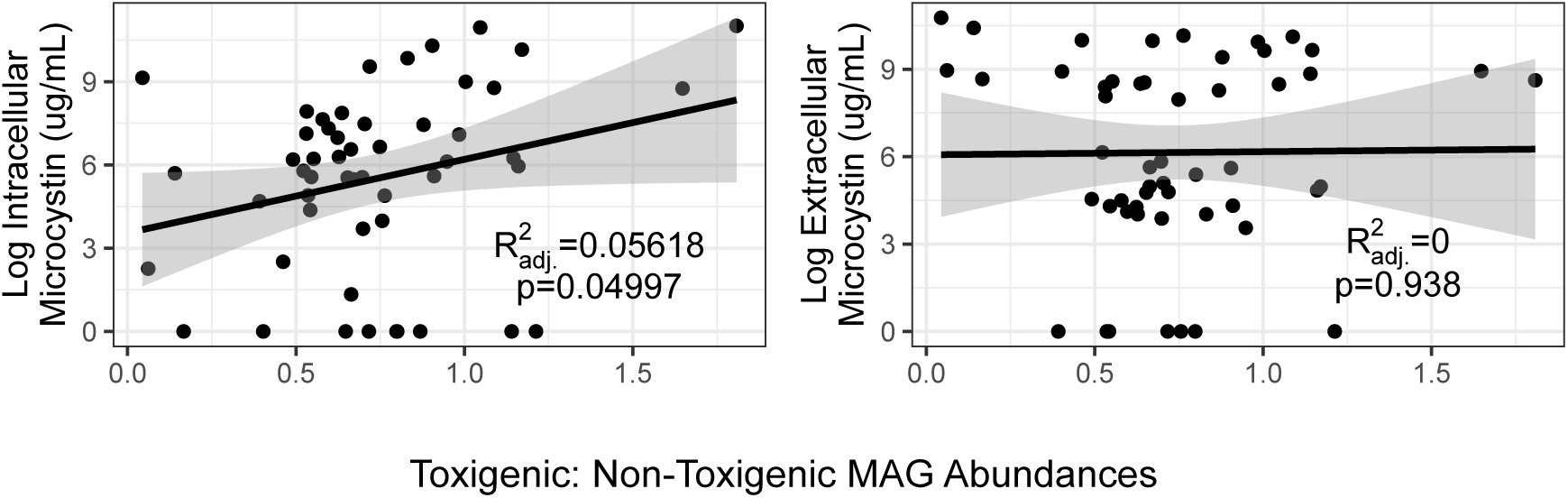
Ratio of toxigenic: non-toxigenic MAGs is weakly associated with intra-cellular, but not extracellular, microcystin concentrations. Correlation between the ratio of toxigenic: non-toxigenic MAGs and log-transformed (left) intracellular and (right) extracellular toxin concentrations. Here, the ambiguous partial producer S53 was assigned to be in the non-toxigenic MAG group. Reported p and *R*^2^ values were extracted from linear regression models run using the respective predictor and response variable.

Moreover, although a statistically significant relationship was identified between intracellular microcystin and the toxigenic: non-toxigenic ratio, this model explained only *↓* 5.6% of the variation in the dataset, suggesting that our models are missing key factors which are important in driving microcystin production and release. In particular, the abundances of key nutrients have shown to be important in predicting microcystin abundances in nature [54], and therefore, models incorporating both nutrient abundances and microbial community metrics may be better suited to describing and predicting intracellular toxin concentrations. One caveat which may further explain the low predictive power of our models is the coarse sampling structure of our study-while we were able to capture variation over nearly the whole year at most sites, monthly sampling may not provide a fine enough resolution to accurately assess the trends in *Microcystis* community composition and toxin release. Notably, microcystin-LA is the most prominent congener in our samples. As this congener can persist in surface waters for weeks, and remain present despite cyanobacterial turnover [55], the abundances of different genomes taken at a given time may not directly reflect the *Microcystis* community composition at the time of toxin release.

While grouping genomes as toxigenic and non-toxigenic is useful for simplifying their possible roles in the community, this lens may be too simplistic to capture the diversity in lifestyles and environmental pressures encountered by different genotypes in our study system. To address this, we conducted multiple linear regression using the abundances of each MAG as an independent variable, and used backwards selection based on AIC to identify the MAGs which best represent the microcystin concentrations.

For intracellular toxin concentrations, the best-fitting model explained 26% of the variation in our dataset, and included toxigenic MAGs S39, S41, S46, and S49 as well as S44, which is ambiguous in its toxin synthesis abilities. Interestingly, the toxigenic MAGs included in this model differed in their direction of association with intracellular toxin concentrations (Figure 5), which might suggest that not all toxigenic genotypes are involved in toxin production in our system.

**Figure 5:**
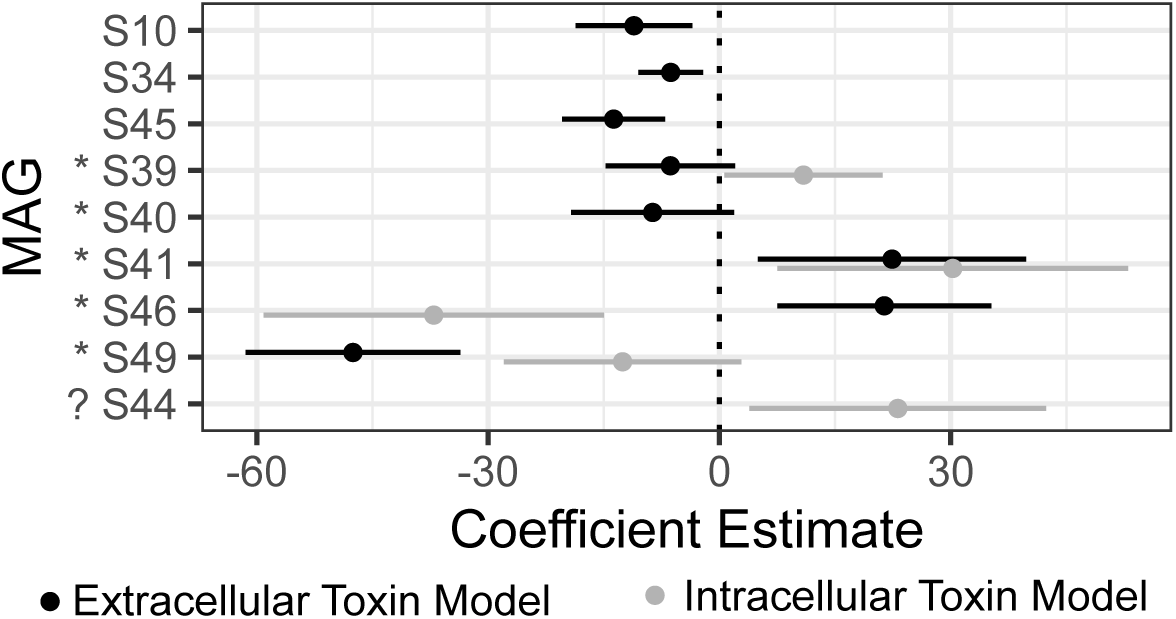
Toxigenic MAGs vary in the directionality of correlation with intracellular and extracellular toxins. Estimated coefficients for MAGs included in the best models for log-transformed intracellular and extracellular toxin concentrations based on AIC. Lines represent 95% confidence intervals for coefficient estimates. MAGs with an asterisk beside their ID are toxigenic. Note that extracellular, but not intracellular, toxin models include non-toxigenic MAGs, and the two models disagree on the direction of impact of some toxigenic MAGs on toxin concentration.

For extracellular toxin concentrations, the best-fitting model explained 64% of the variation in our data and included non-toxigenic MAGs S10, S34, and S45, as well as all of the toxigenic genotypes (Figure 5). While all non-toxigenic MAGs showed negative correlation with extracellular microcystin concentration, toxigenic genomes varied in the direction of their correlation with extracellular toxin concentrations (Figure 5).

The increase in model *R*^2^ values from including the abundances of different MAGs as separate factors might explain why considering only the toxin-producing identity of genomes, through the ratio of toxigenic to non-toxigenic genomes or other metrics, may lead to unexpected results. These results support previous analyses showing that abundances of toxigenic cyanobacteria may not correlate well with toxin concentrations, and suggest that genome- or MAG-level abundances may be better predictors of cyanotoxin production and release [18]. This is further supported by the observation that the direction of association between toxigenic genotype abundances and toxin concentrations may differ between MAGs, suggesting that that microcystin synthesis ability does not necessarily dictate the activities of these microbes in nature. Importantly, some toxigenic genotypes were negatively correlated with toxin abundances, suggesting that some toxigenic MAGs may not be producing cyanotoxins. This highlights the importance of the genotype-level analyses facilitated by metagenomic analysis and supports the idea that considering the abundances of toxigenic genotypes in a specific system is, in many cases, insufficient to predict toxin production.

Microcystin-LA was overwhelmingly dominant at all timepoints in our study system, and therefore, regression models using the total concentration of microcystin toxins as the response variable might more closely resemble the relationships between our microbial communities and microcystin-LA. Therefore, we conducted redundancy analysis, which models each toxin congener separately, to assess whether the abundances of specific MAGs were associated with the concentrations of non-dominant toxin congeners in Valle de Bravo.

The best model for intracellular toxins explained only 16.2% of the variation in our dataset, and included toxigenic MAGs S41, S46, and S49 (Figure 6). Interestingly, all MAGs were anticorrelated with toxin congeners along axis 1, though they showed some correlation with the abundances of the dominant microcystin congener in our system, microcystin LA, along axis 2.

**Figure 6:**
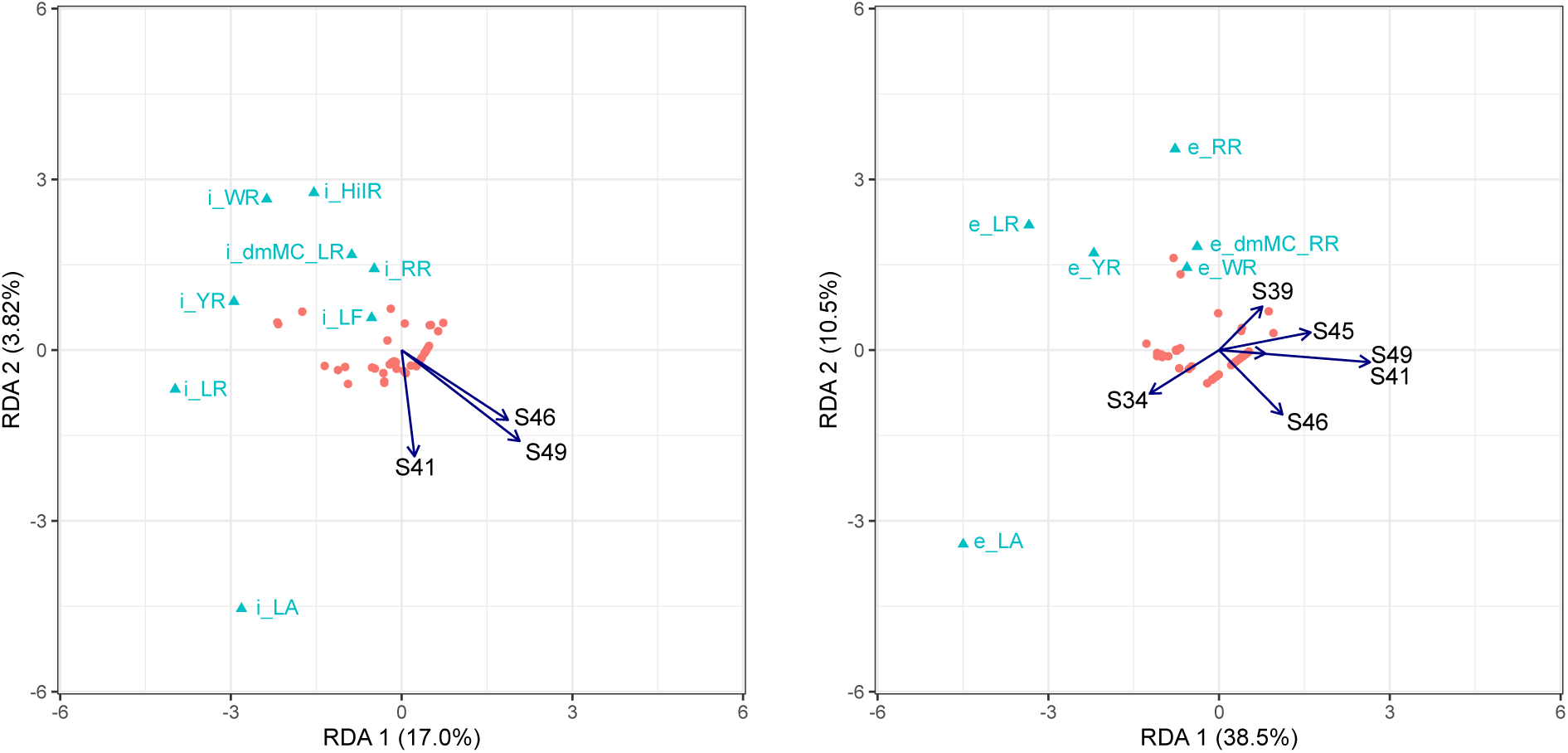
MAG abundances are associated with concentrations of dominant and non-dominant toxin congeners. Biplot of the best fitting models for (left) log transformed intracellular toxin congener concentrations 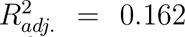 and (right) log-transformed extracellular toxin congener concentrations 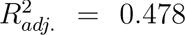, with Hellinger-transformed MAG relative abundances as predictor matrices.

The best-fitting model for extracellular toxin concentrations accounted for a much higher amount of variation in our dataset, with an *R*^2^ of 47.8%. Similar to our intracellular toxin concentration model, we identified negative associations between toxigenic MAGs S41, S46 and S49 with all microcystin congeners along axis 1, and positive correlation with extracellular microcystin LA along axis 2. We also observed correlation between toxigenic MAG S39 and non-toxigenic MAG S45 with non-dominant congeners in our system along axis 2.

Our results indicate that toxigenic MAGs S41, S46 and S49 are negatively associated with non-dominant toxin congeners in both intracellular and extracellular models, and therefore are not likely to be involved in their synthesis or release. However, these MAGs are positively correlated with microcystin-LA along axis 2 in both models, which contrasts our results from the multiple regression analysis where MAGs S46 and S49 were negatively correlated with total toxin concentration. Our results also identify positive correlations between MAGs S39, S45 and non-dominant extracellular toxin congeners, suggesting that these MAGs may be prevalent during periods when non-dominant microcystins are released from cells. This might suggest that MAGs S39 and S45 are primarily involved in community turnover events when non-dominant toxin congeners are released rather than synthesis of these toxins themselves.

### Relationships between microcystin biosynthesis operons and microcystin congener concentrations

MAGs represent a “consensus” genotype and therefore may not capture all the micro diversity present in our communities which may be relevant for production of lower concentration microcystin congeners. Therefore, we mapped our metagenome reads to a reference *mcy* biosynthetic gene cluster from reference genome *Microcystis aeruginosa PCC 7806SL*, which is a well-researched genotype known to produce microcystin-LR. Using these mapped reads, we identified sites of single nucleotide variation (SNV) in the *mcy* biosynthetic gene clusters in our population. We then conducted redundancy analysis to identify nucleotide positions in the *mcy* biosynthetic gene cluster where single nucleotide variation is associated with concentrations of different toxin congeners [19]. We concentrated our analyses of micro diversity within the *mcy* biosynthetic gene cluster to focus on the *mcyA*, *mcyB* and *mcyC* genes, since variation in these regions have been identified in previous studies to affect the form of microcystin produced by a given genotype [12, 56].

Our redundancy analyses identified multiple positions in *mcyABC* where nucleotide variation (the frequency of observing a different nucleotide than what is found in our reference *mcy* biosynthetic gene cluster) is correlated with concentrations of intracellular and extracellular microcystin congeners (Figure 7). These models had relatively high explanatory power, with adjusted *R*^2^ values of around 41% for both intracellular and extracellular toxin models. While many of the SNVs identified represent a difference between our reference strain and the *mcyABC* observed in our MAGs, our analyses identified two positions, *mcyA* 6570 and *mcyB* 3490, where nucleotide substitutions resulted in missense or nonsense mutations. In particular, the SNV at *mcyA* 6570 was an important predictor in the intracellular toxin model, and represents a nonsense mutation introducing an early stop codon resulting in truncation of *mcyA*. Curiously, while it might be expected for this SNV to be negatively correlated with all toxins along both axes, within our ordination plot, this SNV sits close to the center and is slightly positively correlated with all toxin congeners along axis 1.

**Figure 7:**
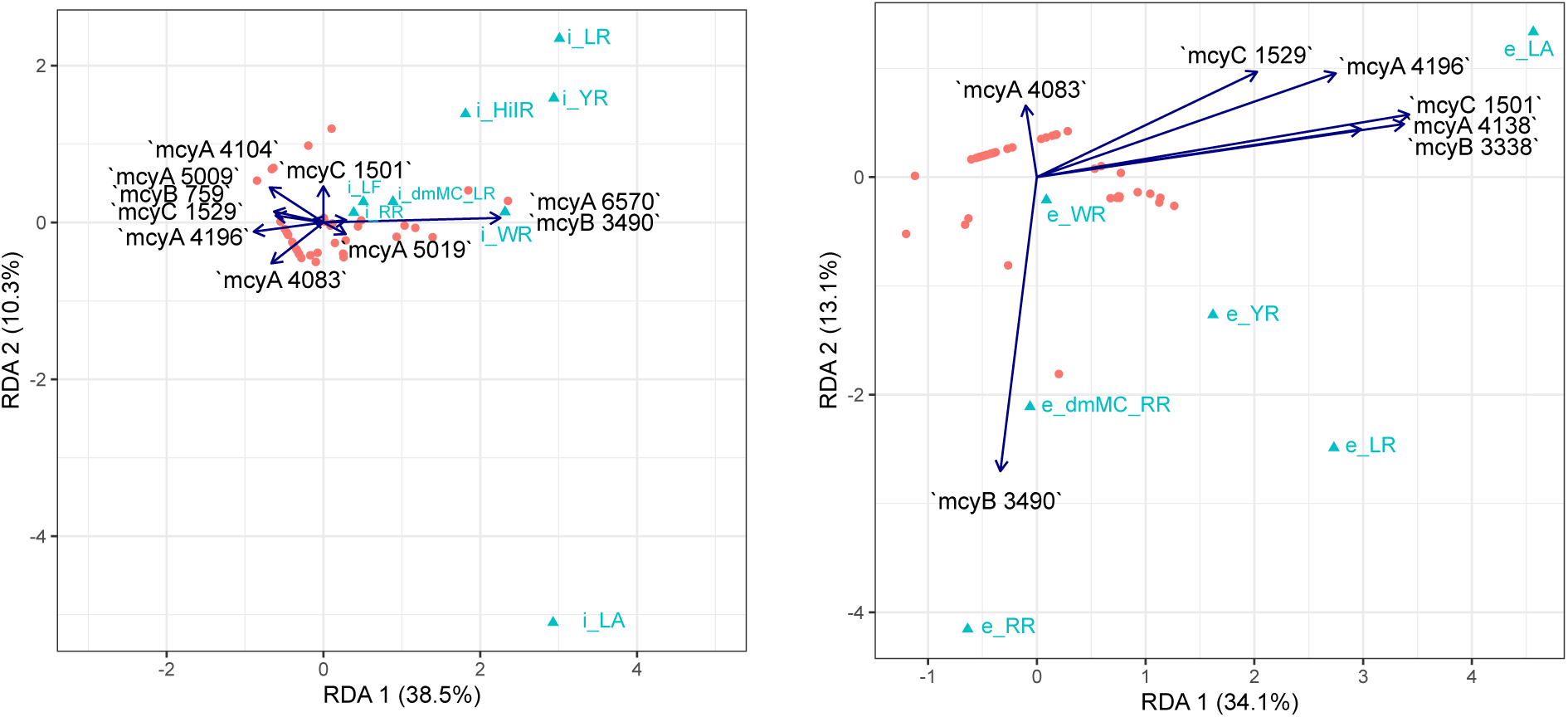
Single nucleotide variation is associated with concentrations of different microcystin congeners. (left) Biplot of the best fitting model of log-transformed intracellular microcystin congener concentrations with SNV frequency within *mcyABC* as predictors. 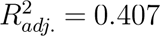 (right) biplot of the best fitting model of log-transformed extracellular microcystin congener concentrations with SNV frequency within *mcyABC* as predictors. 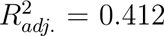

Additionally, the SNV in *mcyB* 3490 corresponds to a missense mutation resulting in a substitution of Proline for Alanine in the synthesized McyB. This SNV is positively correlated with all intracellular microcystin congeners along axis 1. In contrast, this substitution is correlated with only extracellular microcystin RR along axis 1, and all congeners except microcystin-LA along axis 2. These results suggest that substitution at this particular nucleotide, or larger changes in the surrounding region, may be associated with the production of various microcystin congeners, and the release of non-dominant congeners. Previous work has identified that recombination events where a portion of *mcyB* is replaced by genetic material from *mcyC* are responsible for the production of microcystin-RR [56, 12]. As our SNV position falls near the region where these recombination events generally occur, it is possible that the signal detected by our analyses is related to these changes.

Lastly, to determine whether SNV or MAG abundances were better fit to describe the variation in toxin congeners observed in Valle de Bravo, we compared the explanatory power of three redundancy models: one based on MAG abundances (MAG-only), one based on SNV abundances (SNV-only), and one containing both MAG and SNV predictors (combined). These models only included the factors which were identified using backwards AIC model selection to be important in describing the toxin congener community. In both cases, the addition of single nucleotide variation to our models increased the adjusted *R*^2^ values considerably (Figure S5). In the case of intracellular toxin, the SNV predictors appeared to have much more importance than MAG predictors- this is reflected in the high *R*^2^ value for the SNV-only and combination models (both *↓* 40%) as compared to the MAG-only model (*R*^2^ = 17%).

These results highlight the influence of small genetic changes on the production and release of key cyanotoxins in lake environments, as our results indicate that, even with high-resolution MAG-level analysis of community composition, single nucleotide variation may still play a significant role in shaping the observed microcystin congeners.

### Conclusions

Despite these caveats, our results confirm the coexistence of toxigenic and non-toxigenic *Microcystis* genotypes over space and time. The relationship between toxigenicity and toxin abundances is nuanced. First, the relationship differs for intra-and extra-cellular toxins, with a weak but detectable positive correlation between the ratio of toxigenic to non-toxigenic genotypes and intracellular, but not extracellular toxins. Second, the identity of specific genotypes, including both toxigenic and non-toxigenic genotypes, is more informative than relative abundances or ratios of toxigenic to non-toxigenic genotypes in predicting toxin abundances. Finally, single nucleotide variation provides additional insight into the toxin congener abundances, particularly those of non-dominant congeners. Going forward, we hope that this fine-scale genetic perspective on bloom-forming cyanobacteria can be extended to other lakes and cyanobacterial species. It remains to be seen whether adding genotype-level correlates of toxicity, in conjunction with other biotic and abiotic biomarkers, can improve our ability to predict toxic blooms. Nonetheless, we believe that the work presented here demonstrates the utility of genotype-level resolution metagenomic approaches in providing an increased understanding of the factors affecting cyanobacterial bloom formation and cyanotoxin release.

## Data availability

Raw metagenomic sequencing data have been deposited in the NCBI SRA database under BioProject PRJNA1085781. MAGs fasta files are available on Figshare (https://figshare.com/s/9253966b043d791364d5). Other data will be made available on request.

## Acknowledgments

We are thankful to the Mexican Secretaría de Marina, Capitanía de Puerto Valle de Bravo, for the support provided during all the collecting campaigns, including the naval personnel and the use of ships. We thank Julie Marleau and Tahiana Andriananjamanantsoa for participating in DNA extraction and library preparation, and Quoc Tuc Dinh for the analysis of the toxins. Finally, we thank Gavin Douglas for his input and comments that helped improve the paper.

