## Supplemental Tables and Figures for "Linking Cyanobacterial Genomes to Toxin Dynamics Through Genome-Resolved Metagenomics"

### Supplemental Materials

Table S1: Completeness and contamination of our MAGs as assessed by CheckM. Genomes highlighted in orange were removed from our database in this study

| Strain ID | Complete | Contamination |
|---|---|---|
| S53C97.18346NC785 | 76.63 | 1.20 |
| S41C63.18345NC785 | 83.74 | 5.96 |
| S46C30.18339NC785 | 84.60 | 2.33 |
| S16C37.18267NC559 | 86.32 | 7.03 |
| S39C51.18338NC785 | 86.44 | 2.70 |
| S49C42.18343NC785 | 86.67 | 8.84 |
| S44C19.18340NC785 | 86.77 | 2.34 |
| S40C62.18344NC785 | 89.75 | 3.03 |
| S10C14.18282NC559 | 90.63 | 9.17 |
| S48C32.18337NC559 | 98.57 | 6.97 |
| S34C25.25903NC559 | 99.23 | 2.38 |
| S45C20.18335NC559 | 99.45 | 6.54 |
| S50C44.18341NC559 | 92.30 | 14.77 |
| S14C26.18283NC559 | 90.07 | 10.09 |
| S21C62.18286NC785 | 70.43 | 2.08 |
| S17C38.18284NC785 | 54.50 | 1.68 |
| S35C37.25908NC559 | 59.95 | 0.73 |

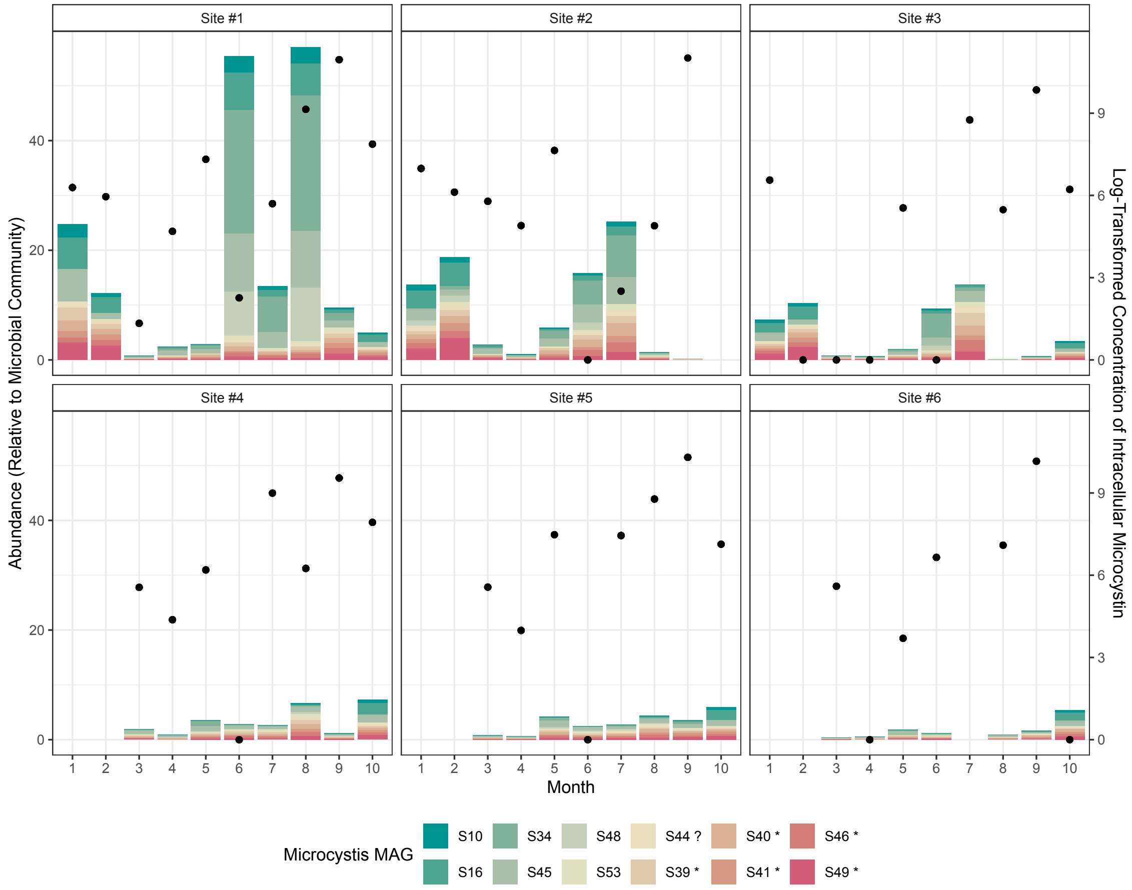

Figure S1: Abundances of *Microcystis* MAG k-mers relative to kmers from all DNA present in our samples at different timepoints and locations in the Valle de Bravo Reservoir. The left y-axis corresponds to the stacked bar plot, where the numerical value represents the proportion of MAG in the community, with MAG IDs included in the legend. MAG IDs with an asterisk beside the name denote toxigenic MAGs (shades of orange/red), while those with a question mark (?) represent ambiguous/ partial microcystin producers. The right y-axis and points represent the log-transformed sum of the concentrations of all intracellular microcystin congeners. Toxigenic and non-toxigenic MAGs co-occurred at all sites and timepoints in our study, and high abundances of microcystin were observed at all sites in June through September. However, increases in toxin concentrations was not always coupled with high abundances of toxigenic MAGs.

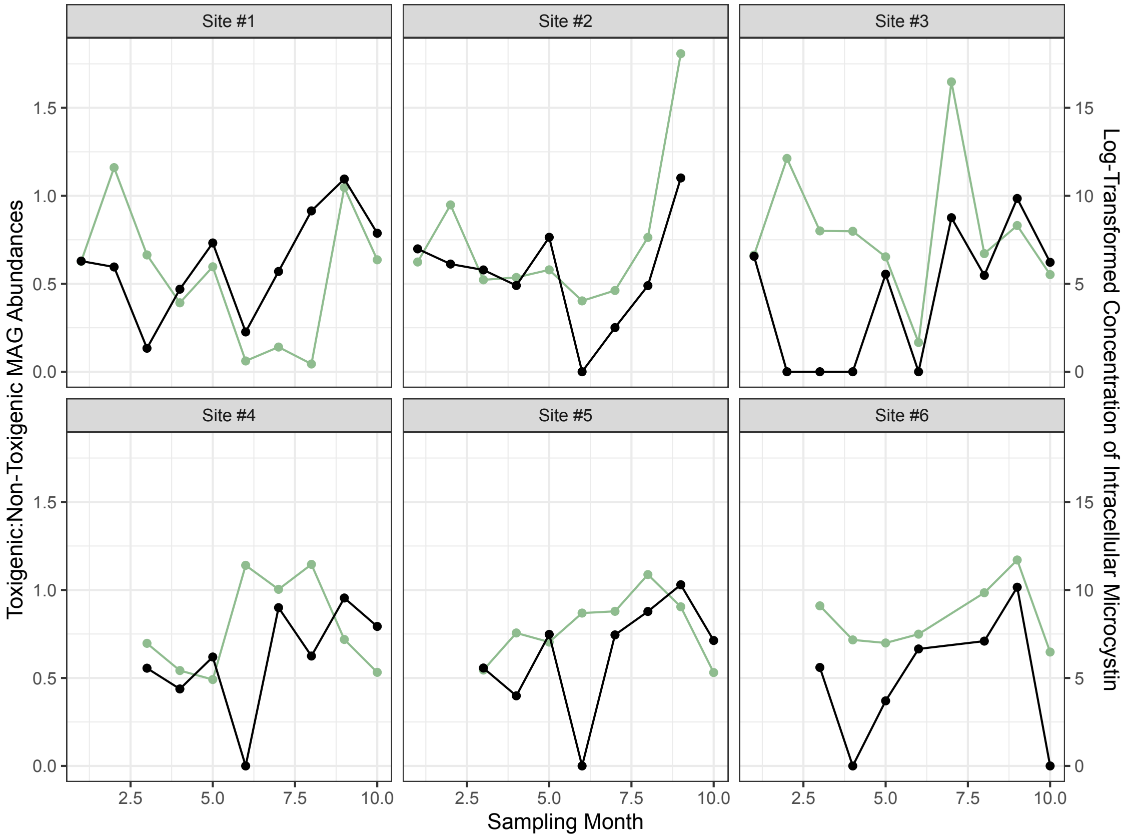

Figure S2: Ratio of toxigenic *Microcystis* MAGs to non-toxigenic MAGs at different timepoints and locations in the Valle de Bravo Reservoir. The left y-axis and green line corresponds to ratio the toxigenic to non-toxigenic MAGs. The right y-axis and black line represents the log-transformed sum of the concentrations of all intracellular microcystin congeners measured in our samples. Here, the ambiguous partial producer S53 was assigned to be in the non-toxigenic MAG group.

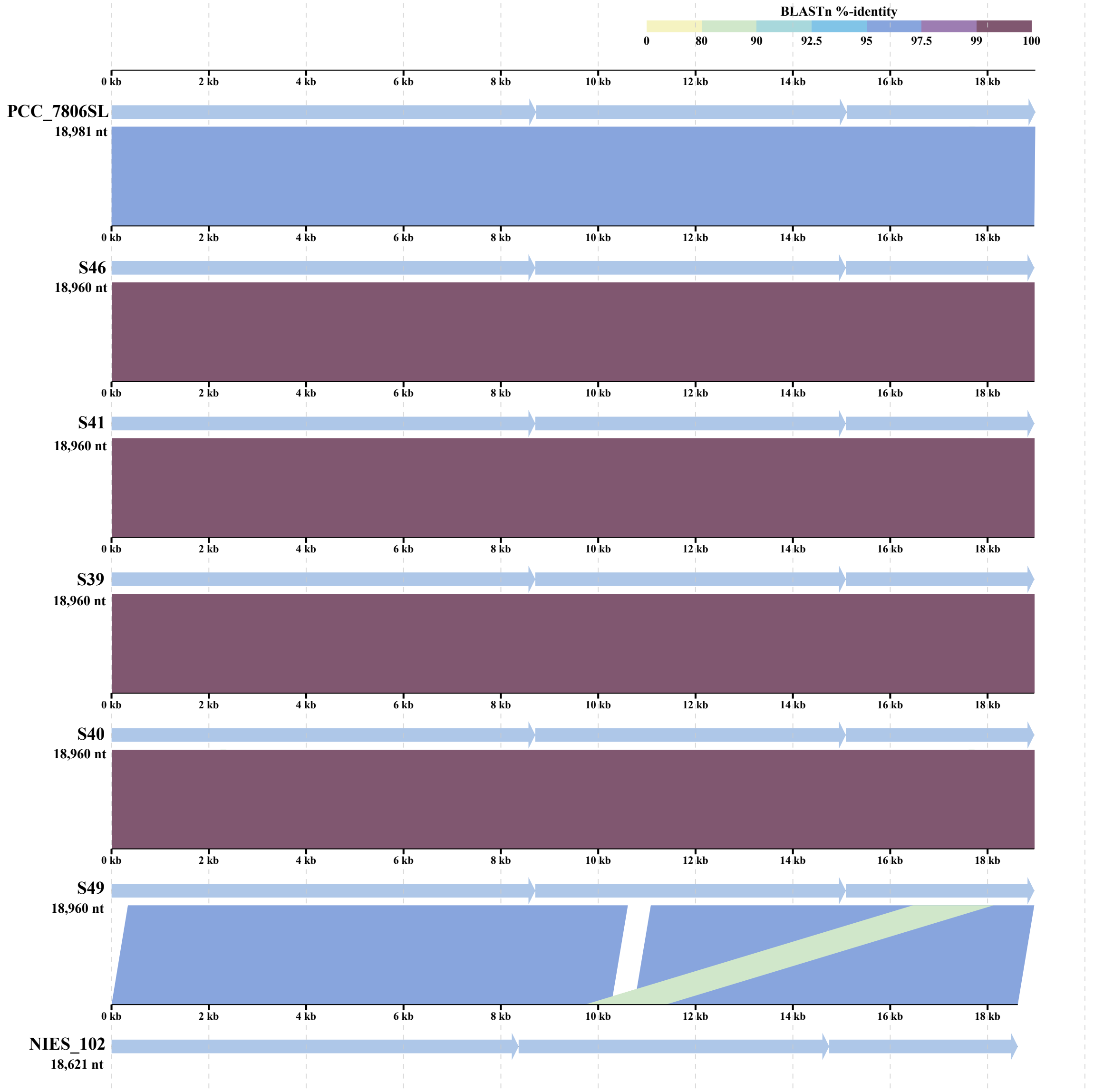

Figure S3: Sequence similarity between the *mcyABC* gene cluster in reference *Microcystis aeruginosa PCC 7806SL* and the assembled MAGs. MAG *mcy* biosynthetic gene clusters are identical, and are highly similar to the reference.

Table S2: BLASTP outputs for *mcy* biosynthetic genes in the *Microcystis* MAGs used in our study

| MAG | Gene | % Identity | Query Coverage | E-Value |
|---|---|---|---|---|
| S39 | mcyJ | 98.2 | 89.4 | 2.70E-165 |
| S39 | mcyI | 96.4 | 100 | 7.30E-185 |
| S39 | mcyH | 95.7 | 92 | 3.50E-291 |
| S39 | mcyG | 95.9 | 100 | 0 |
| S39 | mcyE | 96.3 | 100 | 0 |
| S39 | mcyD | 95.5 | 100 | 0 |
| S39 | mcyA | 94.4 | 96.1 | 0 |
| S39 | mcyB | 96 | 100 | 0 |
| S39 | mcyC | 94 | 100 | 0 |
| S39 | mcyF | 98 | 100 | 4.40E-142 |
| S40 | mcyJ | 98.2 | 89.4 | 2.70E-165 |
| S40 | mcyI | 96.4 | 100 | 7.30E-185 |
| S40 | mcyH | 95.7 | 92 | 3.50E-291 |
| S40 | mcyG | 95.9 | 100 | 0 |
| S40 | mcyE | 96.3 | 100 | 0 |
| S40 | mcyD | 95.5 | 100 | 0 |
| S40 | mcyA | 94.4 | 96.1 | 0 |
| S40 | mcyB | 96 | 100 | 0 |
| S40 | mcyC | 94 | 100 | 0 |
| S40 | mcyF | 98 | 100 | 4.40E-142 |
| S41 | mcyJ | 98.2 | 89.4 | 2.70E-165 |
| S41 | mcyI | 96.4 | 100 | 7.30E-185 |
| S41 | mcyH | 95.7 | 92 | 3.50E-291 |
| S41 | mcyG | 95.9 | 100 | 0 |
| S41 | mcyE | 96.3 | 100 | 0 |
| S41 | mcyD | 95.5 | 100 | 0 |
| S41 | mcyA | 94.4 | 96.1 | 0 |
| S41 | mcyB | 96 | 100 | 0 |
| S41 | mcyC | 94 | 100 | 0 |
| S41 | mcyF | 98 | 100 | 4.40E-142 |
| S44 | mcyI | 96.4 | 100 | 7.30E-185 |
| S44 | mcyH | 95.7 | 92 | 3.50E-291 |
| S44 | mcyG | 95.5 | 100 | 0 |
| S44 | mcyD | 95.9 | 100 | 0 |
| S44 | mcyA | 95 | 100 | 0 |
| S44 | mcyB | 97.4 | 100 | 0 |
| S44 | mcyC | 93.3 | 100 | 0 |
| S44 | mcyF | 98 | 100 | 4.40E-142 |

| | | | | |
|---|---|---|---|---|
| S46 | mcyJ | 98.2 | 89.4 | 2.70E-165 |
| S46 | mcyI | 96.4 | 100 | 7.30E-185 |
| S46 | mcyH | 95.7 | 92 | 3.50E-291 |
| S46 | mcyG | 95.9 | 100 | 0 |
| S46 | mcyE | 96.3 | 100 | 0 |
| S46 | mcyD | 95.5 | 100 | 0 |
| S46 | mcyA | 94.4 | 96.1 | 0 |
| S46 | mcyB | 96 | 100 | 0 |
| S46 | mcyC | 94 | 100 | 0 |
| S46 | mcyF | 98 | 100 | 4.40E-142 |
| S49 | mcyJ | 98.2 | 89.4 | 2.70E-165 |
| S49 | mcyI | 96.4 | 100 | 7.30E-185 |
| S49 | mcyH | 95.7 | 92 | 3.50E-291 |
| S49 | mcyG | 95.9 | 100 | 0 |
| S49 | mcyE | 96.3 | 100 | 0 |
| S49 | mcyD | 95.5 | 100 | 0 |
| S49 | mcyA | 94.4 | 96.1 | 0 |
| S49 | mcyB | 96 | 100 | 0 |
| S49 | mcyC | 94 | 100 | 0 |
| S49 | mcyF | 98 | 100 | 4.40E-142 |
| S53 | mcyI | 96.4 | 100 | 7.30E-185 |
| S53 | mcyH | 95.5 | 92 | 2.20E-290 |
| S53 | mcyC | 93.3 | 100 | 0 |
| S53 | mcyF | 98.3 | 100 | 1.10E-99 |

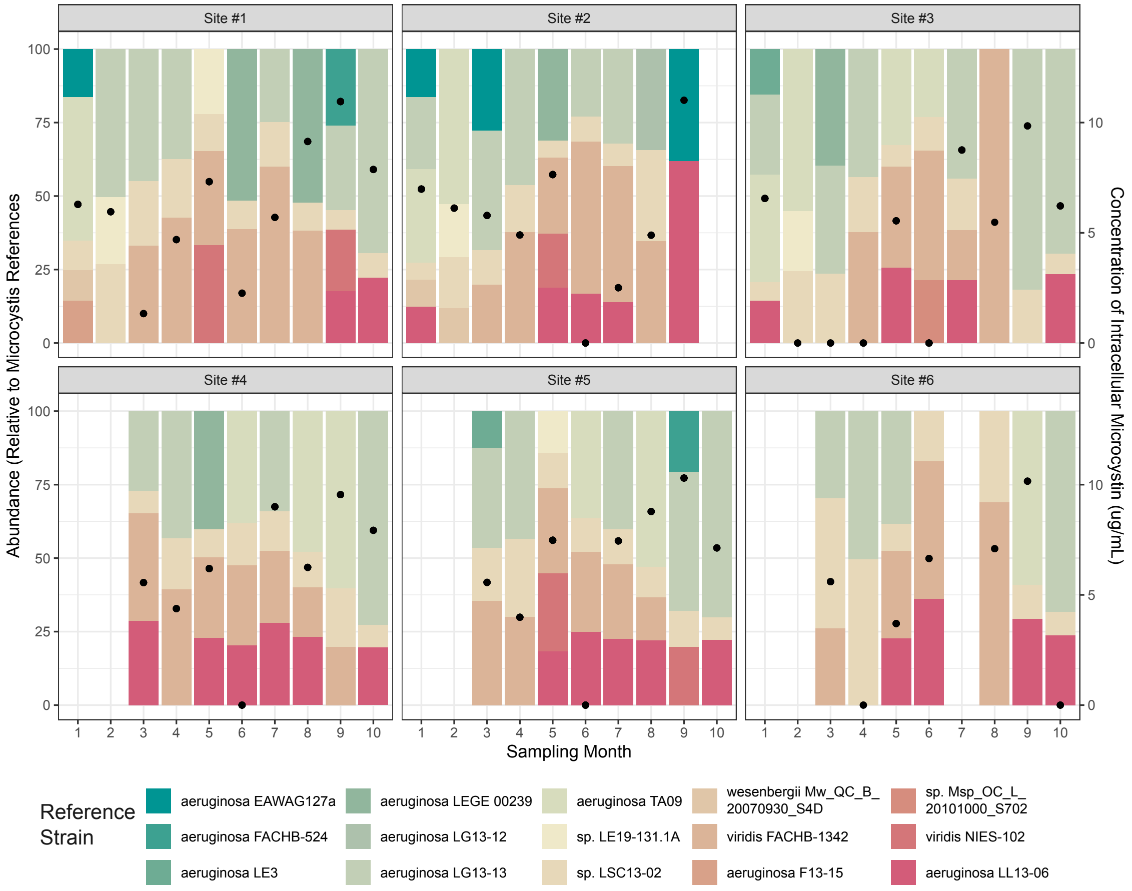

Figure S4: Relative abundances of reference *Microcystis* genotypes downloaded from NCBI at different timepoints and locations in the Valle de Bravo Reservoir. The left y-axis corresponds to the stacked bar plot, where the numerical value represents the relative abundance of the reference genotype in the *Microcystis* community, with genotype names included in the legend. Genotypes with an asterisk beside the name denote toxigenic genomes. The right y-axis and points represent the log-transformed sum of the concentrations of all intracellular microcystin congeners.

| | Parameter | MAG only model | SNV only model | MAG + SNV model |
|---|---|---|---|---|
| anova.cca by margin p-value | S34 | 0.158 | | 0.087 |
| | S39 * | 0.157 | | 0.041 * |
| | S41 * | 0.003 * | | 0.184 |
| | S45 | 0.009 * | | 0.045 * |
| | S46 * | 0.109 | | 0.095 |
| | S49 * | 0.001 * | | 0.003 * |
| | mcyA 4083 | | 0.053 | 0.047 * |
| | mcyA 4183 | | 0.009 * | 0.008 * |
| | mcyA 4196 | | 0.013 * | 0.022 * |
| | mcyB 3338 | | 0.007 * | 0.592 |
| | mcyB 3490 | | 0.002 * | 0.033 |
| | mcyC 1501 | | 0.010 * | 0.329 |
| | mcyC 1529 | | 0.026 * | 0.438 |
| "AIC" | Model | 117.40 | 118.35 | 106.60 |
| Adj R^2 | Model | 0.41 | 0.41 | 0.59 |

| | Parameter | MAG only model | SNV only model | MAG + SNV model |
|---|---|---|---|---|
| anova.cca by margin p-value | S41 * | 0.010 * | | 0.041 * |
| | S46 * | 0.001 * | | 0.648 |
| | S49 * | 0.177 | | 0.156 |
| | mcyA 4083 | | 0.073 | 0.085 |
| | mcyA 4104 | | 0.050 * | 0.043 * |
| | mcyA 4196 | | 0.024 * | 0.066 |
| | mcyA 5009 | | 0.075 | 0.239 |
| | mcyA 5019 | | 0.035 * | 0.094 |
| | mcyA 6570 | | 0.029 * | 0.113 |
| | mcyB 759 | | 0.061 | 0.500 |
| | mcyB 3490 | | 0.019 * | 0.044 * |
| | mcyC 1501 | | 0.045 * | 0.129 |
| | mcyC 1529 | | 0.017 * | 0.051 |
| "AIC" | Model | 141.29 | 132.65 | 131.18 |
| Adj R^2 | Model | 0.17 | 0.41 | 0.45 |

Figure S5: (left) Significance of terms and $R^2$ values for MAG-only, SNV-only, and combined models of log-transformed extracellular toxin concentrations. (right) Significance of terms and $R^2$ values for MAG-only, SNV-only, and combined models of log-transformed intracellular toxin concentrations.
